# G-quadruplex-forming sequences as potential drivers of genetic diversity in primate protein coding genes

**DOI:** 10.1101/2020.08.28.272971

**Authors:** Manuel Jara-Espejo, Sergio Roberto Peres Line

## Abstract

While non-coding G-quadruplexes (G4s) act as conserved regulatory elements when located in gene promoter and splice sites, the G4 evolutionary conservation in protein coding regions have been low explored. To address the evolutionary dynamics acting on coding G4, we mapped and characterized potential G4-forming sequences across twenty-four primate’s gene orthologous. We found that potentially more stable G4 motifs exist in coding regions following a species-specific trend. Moreover, these motifs depicted the least conserved sites across primates at both the DNA and amino acid levels and are characterized by an indel-rich mutational pattern. This trend was not observed for less stable G4 motifs. A deeper analysis revealed that [G_>=3_N_1_]_4_ motifs, depicting potentially most stable G4s, were associated with the lowest conservation and highest indel frequencies. This mutational pattern was more evident when G4-associated amino acid regions were analyzed. We discuss the possibility of an overall conservation of less/moderate stability G4, while more stable G4 may be preserved or arises in a species-specific manner, which may explain their low conservation. Since structure-prone motifs, including G4, have the potential to induce genomic instability, this evolutionary trend may contribute to avoid broad deleterious effects driven by stable G4 on protein function while promoting genetic diversity across close-related species.

## INTRODUCTION

G-quadruplexes (G4) are non-canonical structures formed from guanine-rich sequences in the context of DNA and RNA. The interaction of four guanines through Hoogsteen bonds forms planar G-tetrads, which are stacked to constitute the four-stranded G4 (Sen and Gilbert, 1988; Burge et al, 2006). The G4 is further stabilized by monovalent cations, mainly K+ or Na+, located in the central cavities of the structure (Tateishi-Karimata et al., 2014; Bhattacharyya et al., 2016). DNA G4 have been extensible studied. Computational analyses have demonstrated the enrichment of G4 at important human genome regions as replication origins, telomeric regions, ribosomal DNA, immunoglobulin heavy chain class switch recombination region, and in transcriptional regulatory regions of multiple genes and oncogenes (Maizels and Gray, 2013). Moreover, G4 existence have been demonstrated in vivo (Biffi et al., 2013), and the use of high-throughput sequencing methods has led researchers to experimentally map G4 in human (Chambers et al., 2015) and other organism genomes (Marsico et al., 2019). The distribution of G4s in human chromatin have been investigated using antibody-based G4 chromatin immunoprecipitation (Hänsel-Hertsch et al., 2016; Hänsel-Hertsch et al., 2018). These studies demonstrated that ∼10000 regions in the human genome can effectively fold into G4s; moreover, these G4 structures are strongly enriched at gene promoters and are strongly associated with elevated transcriptional activity.

RNA G4 (rG4) have been computationally mapped within human genes and are enriched at the TSS (Huppert & Balasubramanian S, 2007), the 5′-UTR (Huppert et al., 2008), and the 5′ end of the first intron (Eddy and Maizels, 2008) and depleted in coding regions (Maizels and Gray, 2013). A recent study demonstrated that stable G4 are strongly depleted from the human coding genome fraction through codon bias selection (Mirihana Arachchilage et al..2019).The deletion of coding G4 may be due to their negative influence on translation (Endoh et al., 2013; Endoh and Sugimoto, 2016; Shabalina et al., 2006; Benhalevy et al., 2017). The genome-wide underrepresentation of stable G4 motifs (G3N1)4, where N is any nucleotide, seems to follow trends depending on the phylogenetic group analyzed; while thermodynamically very stable G4 motifs having identical G, C or T single loop composition tend to be suppressed, the (G3A1)4 motif is strongly preserved in mammals, especially in primates (Puig Lombardi et al., 2019). Moreover, the authors demonstrated that elevated genetic instability in coding regions is directly associated with high G4 stability. The pattern of conservation of stable G4 motifs suggests the maintenance of positive G4-associated biological roles while potential deleterious effects are eliminated during evolution.

Several studies have shown that the occurrence of stable G4 at regulatory non-coding regions, mainly gene promoters, is conserved across several organisms (Rawal et al., 2006; Capra et al., 2010; Marsico et al., 2019). The approaches used are based on the identification through of potential G4-forming sequences (PQS) using computational and/or DNA sequencing methods in different genomes, with a posterior comparison of the pattern of PQS co-occurrence in multiple species at specific genome locations. The result is a well characterized map of non-coding PQSs across species. Within coding regions, evolutionary forces act both against (Eyrewalker & Bulmer, 1993; Kudla et al., 2009) and in favor (Katz and Burge, 2003) of stable RNA secondary structures. G4 tend to be more stable in RNA when compared to their DNA counterparts (Joachimi et al., 2009; Arora and Maiti, 2009). Studies have shown that stable G4 structure present within the CDS can cause ribosomes stalling and that protein expression can be enhanced through silent mutations affecting the G4 stability (Agarwala et al., 2015; Endoh et al., 2013). This suggest functional implications derived from G4 folding/unfolding dynamics in coding regions. However, the characterization of the G4 evolutionary conservation within coding regions has received low attention, likely based on the know G4 broad depletion within exons (Maizels & Gray, 2013).

Here, we report a comprehensive genome-wide analysis of G4 evolutionary trends within protein coding regions. Based on the highest conservation among primate genomes, we selected these species for our study, focusing on PQS location and exact sequence conservation analysis. Although *in silico* PQS identification based on canonical G4 motif (G_3_N_1-7_G_3_N_1-7_G_3_N_1-7_G_3_) can overestimate the number of genomic PQS, these algorithms offer valuable tools to characterize potential G4 structures (Kikin et al., 2006; Huppert & Balasubramanian, 2007). Moreover, the use of motif based PQS identification can be complemented with the estimation of G4 folding energies (Lorenz et al., 2013) in order to have a more realistic view of PQS folding potential. Experimental validation of PQS occurrence and the correlation of these results with previous *in silico* PQS mapping highlight their usefulness (Hänsel-Hertsch et al., 2016; Hänsel-Hertsch et al., 2018). Our analysis was not restricted to the mapping and comparison of human-specific coding G4; all PQS were identified and analyzed irrespective of the primate genome of origin.

We found that PQS stability is strongly associated with their low frequency and conservation among primates. Multiple species genome comparison revealed that more stable PQS tend to occur in single or close-related primates. High PQS stability correlated with a marked drop of conservation at both the DNA and amino acid levels. In contrast, less/moderate stable PQSs are mostly conserved. Strikingly, the highest influence on conservation was observed for the most stable PQS (G_3_N_1_G_3_N_1_G_3_N_1_G_3_). A deeper analysis of the G4-associated mutational pattern in CDSs show that low conservation is supported by an indel enrichment at more stable PQS sites. Overall, we provided evidence that potentially stable G4 in coding regions are evolutionary less conserved among primates. Thus, our study provides insight into the role of G4 structures as potential drivers of genetic diversity in the protein-coding regions of primates, contributing to our understanding of the evolutionary dynamics controlling the evolution of stable DNA/RNA structures.

## RESULTS

### Mapping Potential G4 forming sequences across orthologous primate genes

To assess the genome-wide occurrence of potential G4 sequences in coding regions, we obtained, processed and aligned ortholog CDS sequences from twenty-four primates (see Table 1). PQSs of the form G_3_+L_≤7_G_3+_L_≤7_G_3+_L_≤7_G_3+_ were mapped across 17780 multiple species alignments. Although we used a human-based orthology, using previously aligned sequences let us to perform a non-human biased PQS search in primate orthologous genes. The number of PQSs varied across species (see Fig. 1A); rhesus macaque (n = 1119) and chimpanzee (n = 1082) had the highest PQS prevalence, while bushbaby (n = 455) and tarsier (n = 349) had the lowest number of identified PQSs. Overall, 2937 primate genes had at least one PQS. The PQSs count was strongly correlated with the available orthologous across primates (Spearman’s *rho* = 0.81, p = 4.621e^−0.6^) and moderately correlated with phylogenetic distances to human (Spearman’s *rho* = −0.41, p = 0.047). Some primate genome assemblies, particularly mouse lemur and bushbaby, are largely incomplete. Thus, the PQS count variability is likely a consequence of assembly quality.

**Table 1.**
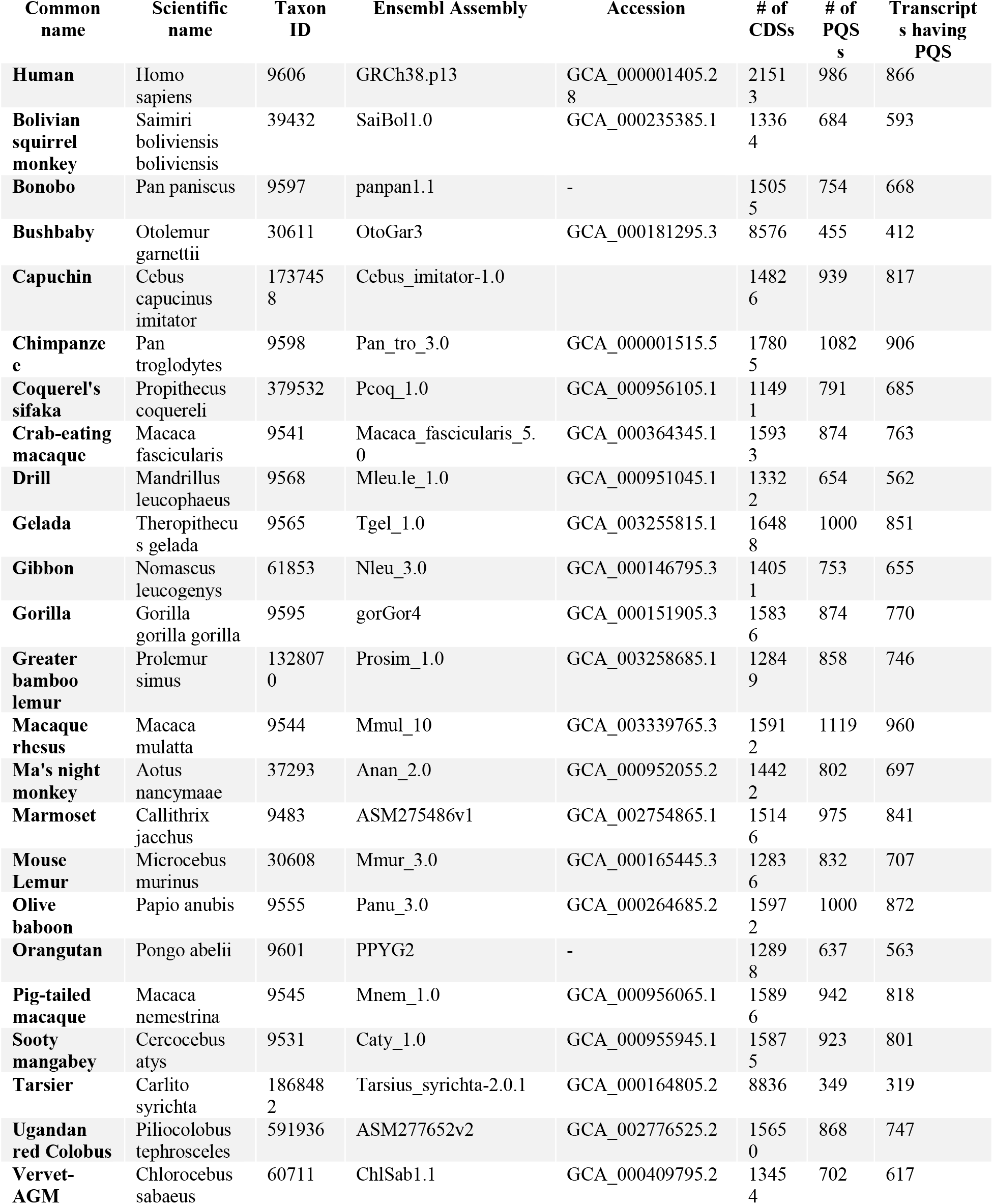
Genomic data sources and PQS count for all analyzed species.

**Fig. 1.**
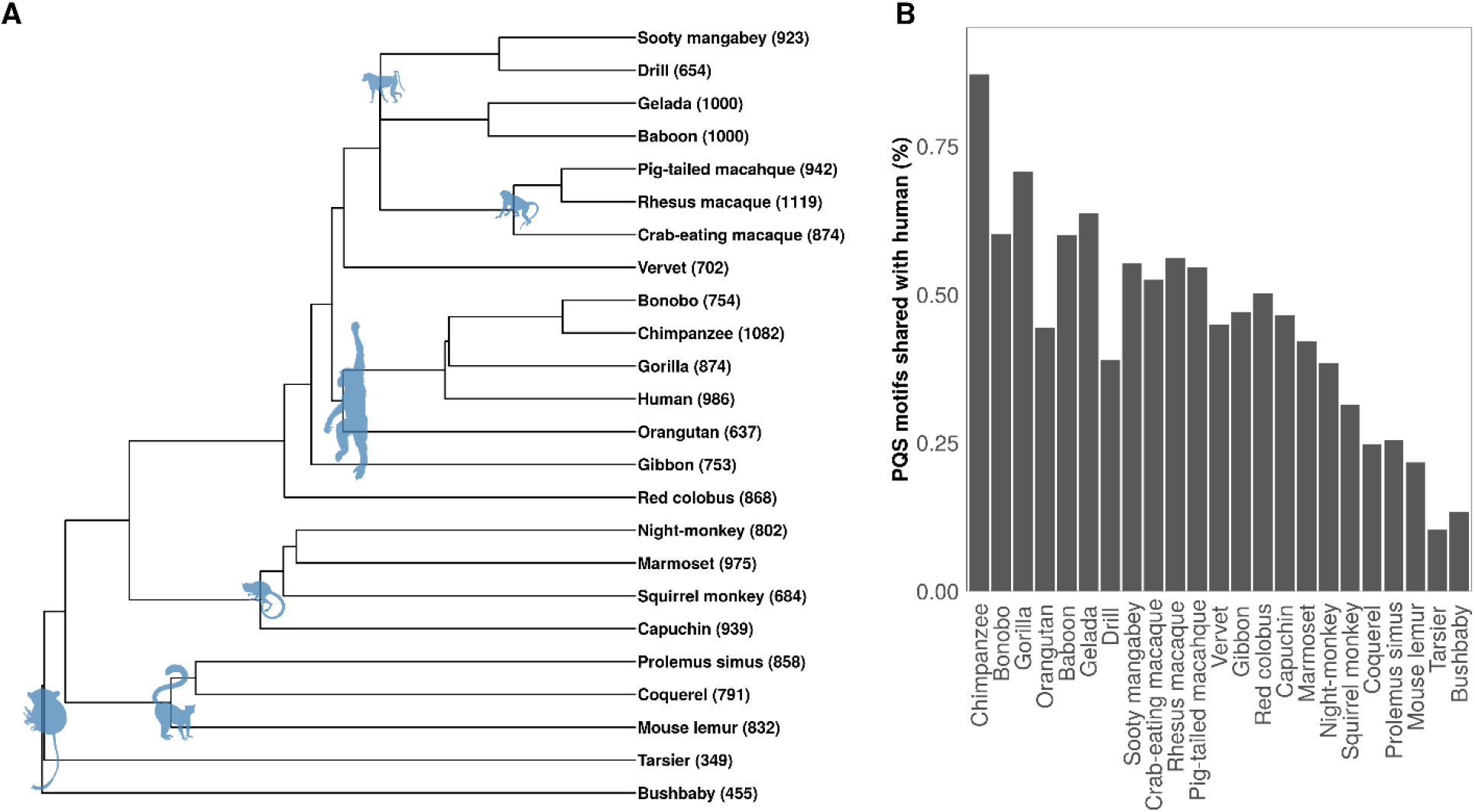
**A.** Phylogenetic representation of 24 primate species analyzed. Total count of PQS of the form G_3+_L_≤7_G_3+_L_≤7_G_3+_L_≤7_G_3+_ identified for each species is indicated in brackets. **B.** Human-centric pairwise comparisons for shared PQS motifs across 17780 orthologous genes.

We analyzed the influence of phylogenetic relationship on PQS occurrences by estimating the fraction of PQS co-occurrence (PQSs in ≥2 species). Using a human-centric (930 PQSs) species × species comparison, we found that the fraction of PQS co-occurrence strongly reflected the phylogenetic distance between human and other species (Spearman’s *rho* = −0.88, p = 2.57e-0.8; Fig. 1B). PQS co-occurrence fractions were 87 %, 45% and 10% for human × chimpanzee, human × vervet and human × tarsier comparisons, respectively. Together, PQS co-occurrence analysis results suggest a high degree of PQSs co-localization between closely related primate species.

### High stability PQSs mark low conserved coding regions across primates

To assess the effect of potential G4 sequences on orthologous genes conservation, DNA PQS conservation was analyzed using a sliding window approach based on their stability. We found that CDS conservation scores are negatively correlated with PQS stability (Fig. 2A). While LS-PQSs (Mfe ≥ −10 kcal/mol, n = 4648) mapped highly conserved regions, HS-PQSs (Mfe ≤ - 30kcal/mol, n = 366) are associated with the least conserved regions within CDSs. Moreover, we noted a very low conservation variability between PQSs and their flanking regions (Fig. 2B). We used the averaged full-length CDSs conservation (CDS_ConScore_) to compute PQSs Z-scores (i.e. effect sizes; see methods) and compare them based on motif predicted stability. We found that PQS and flanking regions conservation varied depending on predicted stability. Overall CDS conservation mostly matched that of LS-PQS; conversely, HS-PQSs exhibited a strong trend to be less conserved than CDSs. Indeed, when compared to fragments of the same length, HS-PQSs were the least conserved region in 60% of the CDS, while LS-PQSs was the least conserved region in only 9% (Fisher’s exact test *p < 2.2*e^−16^, odds ratio = 14.86). These results indicated that more stable PQS motifs and their flanking regions depict clusters of low nucleotide conservation when primate orthologous genes are compared.

**Fig. 2.**
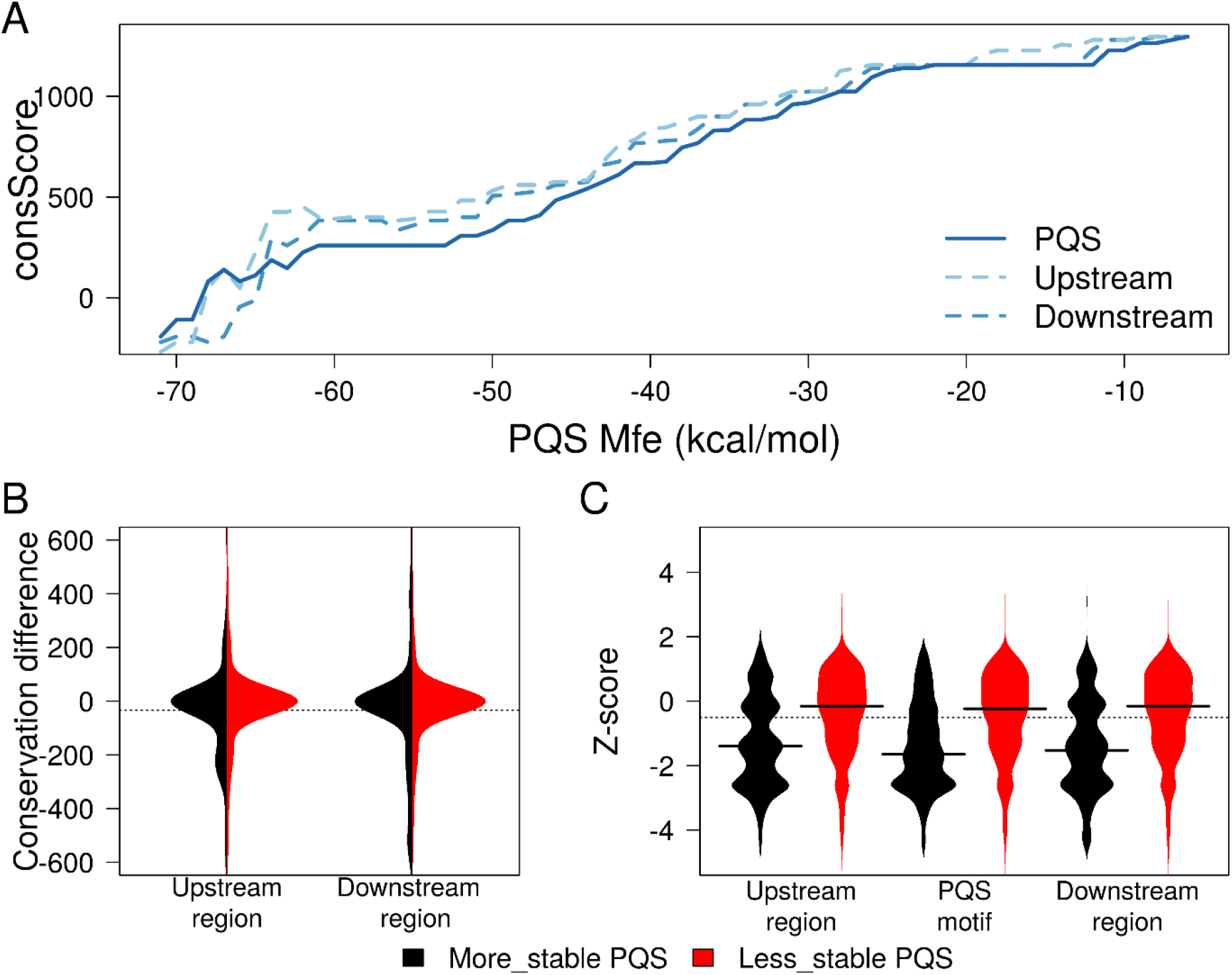
Analysis of DNA conservation at PQS motifs. **A.** Note that as higher the predicted PQS stability, lower the conservation. In addition, conservation of PQS flanking regions is also plotted based on the stability of their PQSs. **B.** Beanplot of conservation variability between PQSs and their flanking regions. **C.** Beanplot of Z-score conservation for PQSs and flanking regions. Lower Z-score values indicate lack of conservation.

Since other than PQS short and stable motifs must be abundant in cDNA, we tested if coding DNA low conservation is intrinsically associated to structurally stable regions (Fig 3. Central panel). We found an overall low conservation pattern associated to more stable random regions (HS-Rand) when compared to those less stable (LS-Rand) (Wilcoxon’s Rank Sum Test p = 8.31 e^−06^). While LS-PQSs and LS-Rand motifs were mostly well conserved, HS-PQSs were significantly less conserved than HS-Rand (Wilcoxon’s Rank Sum Test p = 6e^−09^) and depicted the least conserved group (Fig 3, lower panel). It is important to note that low conserved PQS and random regions tended to be underrepresented within CDSs. These results suggest that cDNA regions with high predicted stability are associated with low nucleotide conservation and that this pattern is more evident if the region can fold into a G4 structure.

**Fig. 3.**
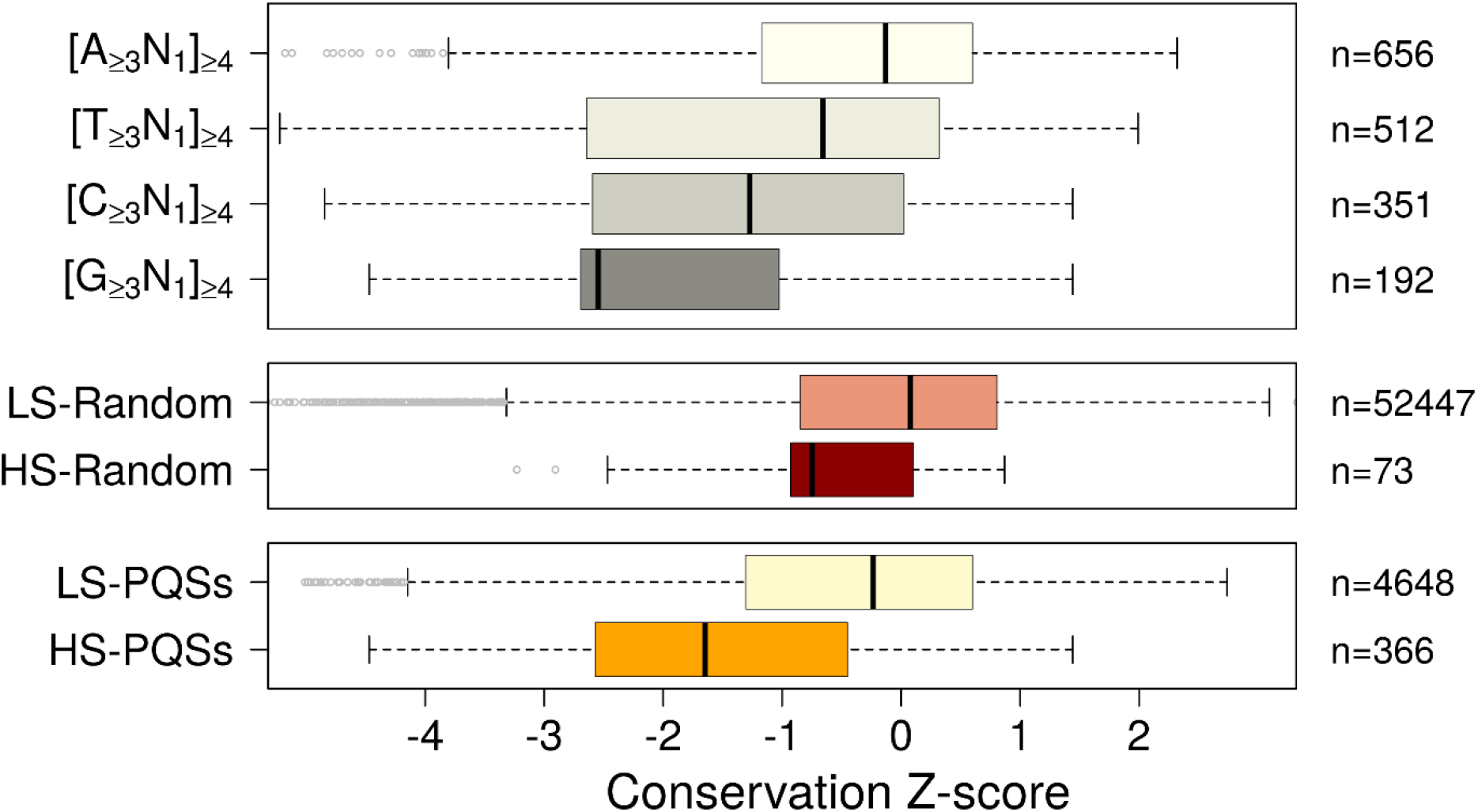
Conservation comparison between PQSs (lower panel), random structured regions (middle) and tandem repeat (upper panel) motifs. Middle panel shows the conservation from low stability (LS-Random) and high stability (HS-Random) non-PQS random regions. Upper panel shows the conservation of A, T, C and G-rich tandem repeats. Note that high stability G-quadruplexes (HS-PQSs and [G>=3N1]]4) are the least conserved regions.

Short tandem repeats can induce mutations within coding and non-coding genome regions (Shah and Mirkin, 2015). So, we asked if the lack of conservation we observed associated to PQSs can also be present around similar tandem regions. We compared the conservation z-score of 5′-N_x_L_1_N_x_L_1_N_x_L_1_N_x_-3′ motifs, where N can be A, T, C or G, x ≥ 3 nt and L represents single nucleotide loops. We found an increasing A < T < C < G lack of DNA conservation inversely correlated to the number of motifs identified. AT-rich tandem regions (n = 1168) appeared to be more conserved. C-rich tandem regions are markedly low conserved, like HS-PQSs (Pairwise Wilcoxon’s Rank Sum Test with P value adjustment for multiple comparisons, *P = 0.18*). Interestingly, G-rich tandem regions, potentially folded into the most stable PQSs, were the least conserved regions, even when compared to HS-PQSs (Pairwise Wilcoxon’s Rank Sum Test with P value adjustment for multiple comparisons, *P* < 2.1e^−05^. Table 2). These regions were less frequent than A, G and C tandem repeats (Fig. 3, upper panel). Thus, G-rich regions potentially folding into highly stable G4s mark low conserved regions and seems to be evolutionary unfavored in CDSs.

**Table 2.** Results from DNA conservation pairwise comparisons between groups

|  | HS-PQS | LS-PQS | HS-Rand | LS-Rand | [G<sub>>=3</sub>N<sub>1</sub>]<sub>4</sub> | [C<sub>>=3</sub>N<sub>1</sub>]<sub>4</sub> | [T<sub>>=3</sub>N<sub>1</sub>]<sub>4</sub> |
| --- | --- | --- | --- | --- | --- | --- | --- |
| **LS-PQS** | $< 2.1e^{-16}$ | - | - | - | - | - | - |
| **HS-Rand** | $3.4e^{-07}$ | 1 | - | - | - | - | - |
| **LS-Rand** | $3.4e^{-07}$ | 1 | 1 | - | - | - | - |
| **[G<sub>>=3</sub>N<sub>1</sub>]<sub>4</sub>** | $2.1e^{-05}$ | $< 2.1e^{-16}$ | $2.5e^{-12}$ | $2.5e^{-12}$ | - | - | - |
| **[C<sub>>=3</sub>N<sub>1</sub>]<sub>4</sub>** | 0.18 | $< 2.1e^{-16}$ | 0.0078 | 0.0078 | $9e^{-06}$ | - | - |
| **[T<sub>>=3</sub>N<sub>1</sub>]<sub>4</sub>** | 0.0104 | $9.4e^{-13}$ | 1 | 1 | $1e^{-08}$ | - | - |
| **[A<sub>>=3</sub>N<sub>1</sub>]<sub>4</sub>** | $< 2.1e^{-16}$ | 1 | 0.2330 | 0.011 | $2.1e^{-16}$ | $2.1e^{-16}$ | $3e^{-10}$ |

*p.adj value using Bonferroni method*

Using the same approach previously described for nucleotide-based conservation analysis, we assessed amino acid (AA) conservation at PQS motifs. We found that PQS-associated AAs (PQS-AAs) exhibited a marked drop of conservation as PQS stability increased (Fig 4A). HS-PQSs, but not LS-PQSs, are significantly associated to the lack of AA conservation (Wilcoxon’s Rank Sum Test, *p* < 2.1e^−16^) across orthologs. Moreover, random non-PQSs regions (n = 10 for each PQS) exhibited a more conserved pattern, like that of LS-PQSs (Fig. 4B).

**Fig. 4.**
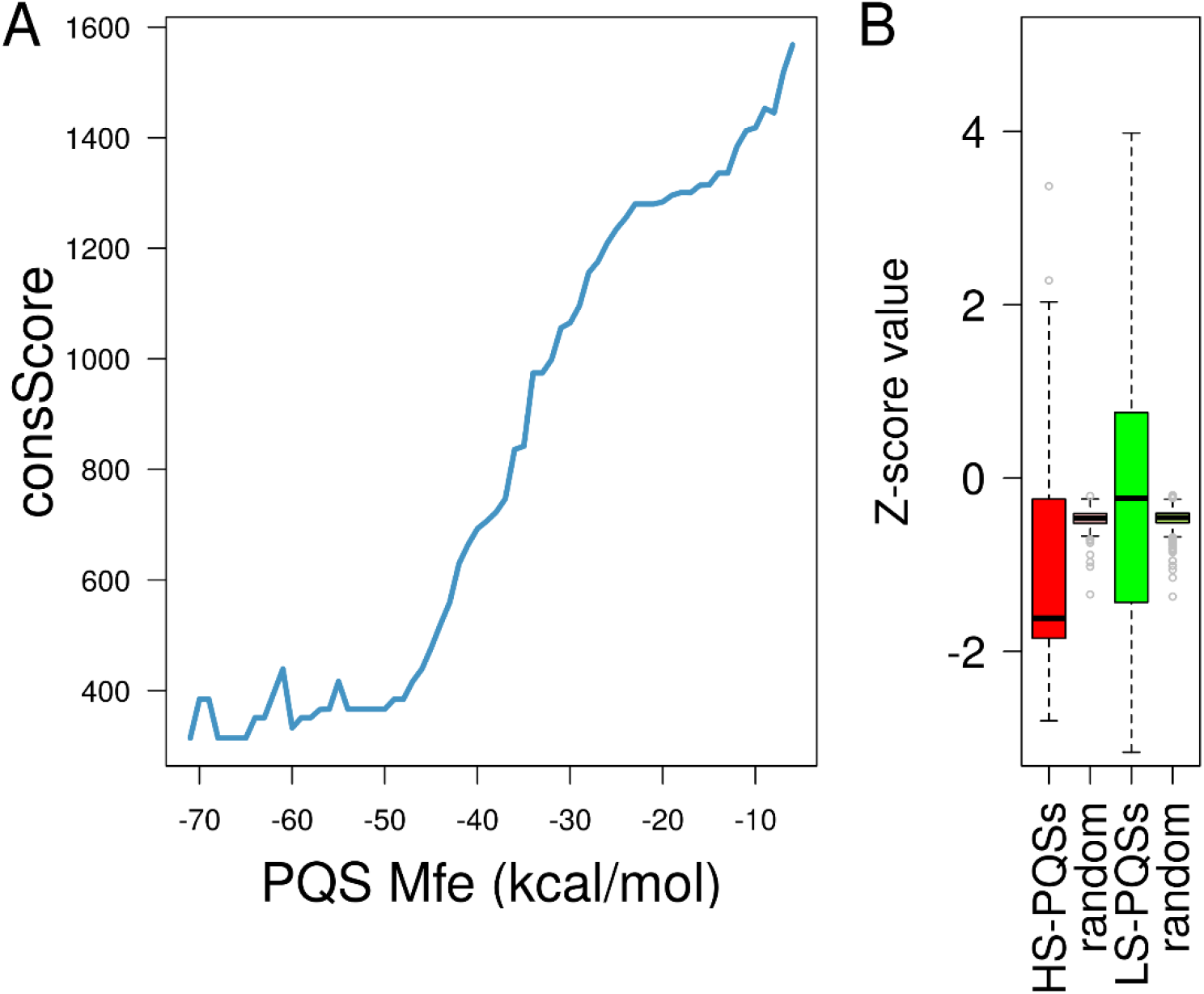
**A** Amino acid conservation comparison between PQSs and random coding regions. Note that conservation decreased and remained low as the PQSs stability increased (∼-30). **B** More stable PQSs marked the least conserved coding regions at the AA level.

Our results indicate that sequence homology among primates seems to be impaired by PQSs presence at both the DNA and amino acid levels; moreover, this lack of conservation strongly depends on PQS structural stability.

### Coding regions rich in indels overlap high stability G4 motifs in primates

The above results indicate that occurrence of HS-PQS is associated with lack of conservation across orthologous in primates. To clarify the mutagenic pattern associated to HS-PQSs, we calculated DNA substitution and indel scores for PQSs and tandem repeat motifs. All analyzed motifs exhibited overall low substitution scores, indicating that low conservation is not a consequence of nucleotide similarity lack (Fig. 5A). Conversely, indel rates varied among motifs; higher values indicated gap-rich consensus sequence, i.e. insertions are favored (Fig. 5B). HS-PQSs indel rates were higher than LS-PQSs (0.19 and 0.10, respectively; Pairwise Wilcoxon’s Rank Sum Test with P value adjustment for multiple comparisons, *P* < 2e^−16^). For tandem repeats we found a growing pattern A < T < C << G of indel rates. G-rich tandem regions exhibited the highest scores when compared to all groups analyzed, including HS-PQSs (0.55 and 0.19, respectively; Pairwise Wilcoxon’s Rank Sum Test with P value adjustment for multiple comparisons, *P < 7*.2e^−07^; Table 3). Half of total PQSs mapped across CDSs (57%) depicted unique G4 motifs, i.e. they were mapped as the G4 sequence with highest potential for one species. Interestingly, when PQS stability was considered, HS-PQSs were significantly more species-specific than LS-PQSs (71% and 53%, respectively; Fisher’s exact test *p*= 6.5e^−16^, odds ratio = 2.05). These results indicate that more stable PQSs may be promoting lack of nucleotide conservation by inducing base insertion mutations in specific- or closely related species.

**Table 3.** Results from DNA indel rates pairwise comparisons between groups

|  | HS-PQS | LS-PQS | [G<sub>>=3</sub>N<sub>1</sub>]<sub>4</sub> | [C<sub>>=3</sub>N<sub>1</sub>]<sub>4</sub> | [T<sub>>=3</sub>N<sub>1</sub>]<sub>4</sub> |
| --- | --- | --- | --- | --- | --- |
| **LS-PQS** | $< 2.1e^{-16}$ | - | - | - | - |
| **[G<sub>>=3</sub>N<sub>1</sub>]<sub>4</sub>** | $2.1e^{-06}$ | $< 2.1e^{-16}$ | - | - | - |
| **[C<sub>>=3</sub>N<sub>1</sub>]<sub>4</sub>** | 0.24 | $< 2.1e^{-16}$ | $1.7e^{-10}$ | - | - |
| **[T<sub>>=3</sub>N<sub>1</sub>]<sub>4</sub>** | 0.00067 | $< 2.1e^{-16}$ | $1.1e^{-13}$ | 1 | - |
| **[A<sub>>=3</sub>N<sub>1</sub>]<sub>4</sub>** | $< 2.1e^{-16}$ | 0.00089 | $< 2.1e^{-16}$ | $1.9.1e^{-12}$ | $6.6e^{-09}$ |

*p.adj value using Bonferroni method*

**Fig. 5.**
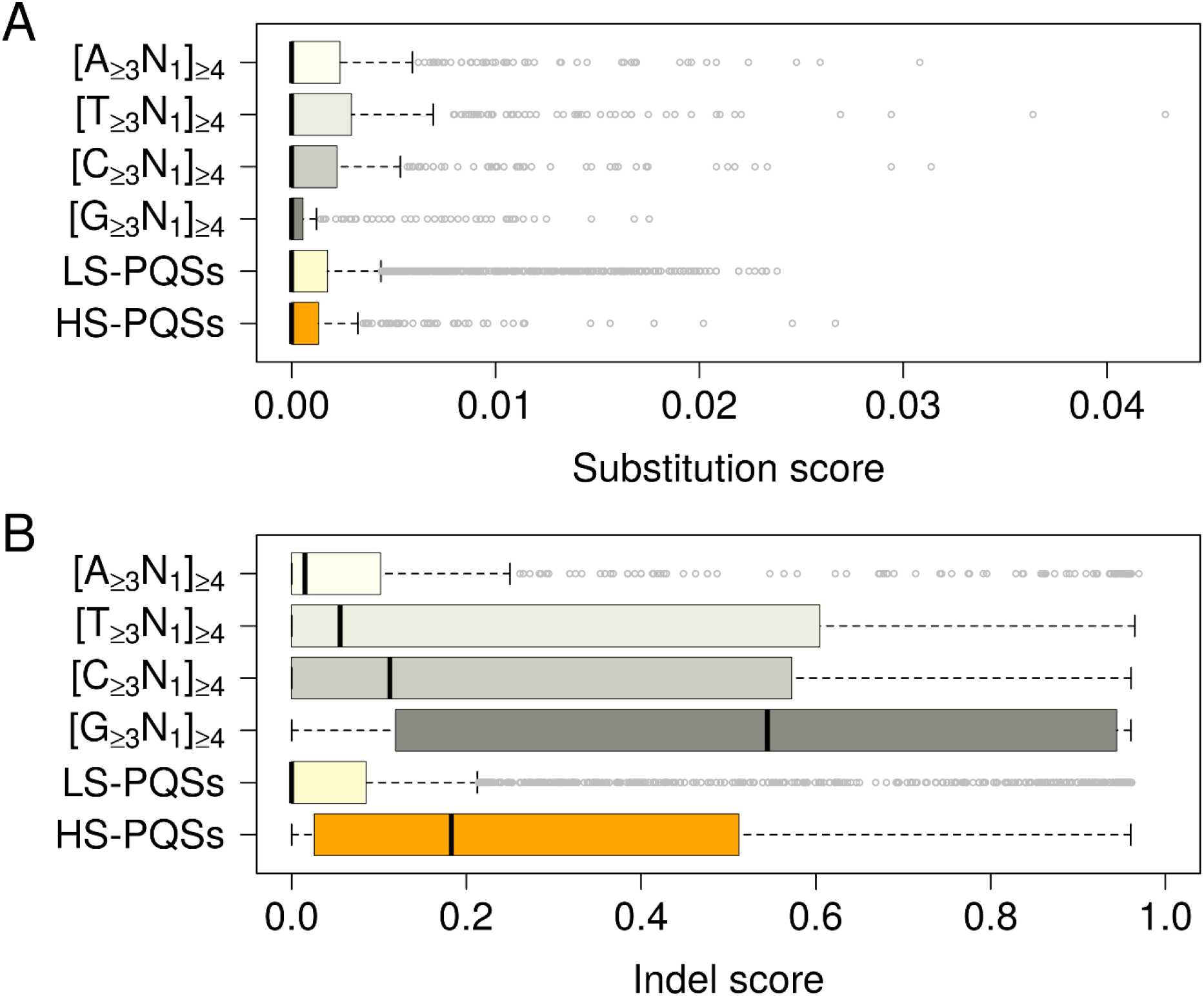
DNA mutational rates for PQSs and tandem repeat regions. **A** Substitution score analysis show that all analyzed motifs exhibited low lack of DNA similarity across orthologous genes. **B** Indel score analysis, higher indel rates are associated to potential structurally stable G4 motifs.

Since alignments are generally more accurate when based on amino acids than on their corresponding nucleotides (Abascal et al., 2010), we additionally assessed the mutational pattern for PQS and tandem repeat motifs at the amino acid level. We observed a slight substitution rate increase for all motifs at the AA level, but not statistical differences were observed (Fig. 6A). Conversely, HS-PQSs showed evident higher indel rates than LS-PQSs (0.51 and 0.03, respectively; Pairwise Wilcoxon’s Rank Sum Test with P value adjustment for multiple comparisons, *P* < 2e^−16^) (Fig 6B). Strikingly, G-rich tandem motifs exhibited the highest AA indel rates (median = 0.86) among all motifs; moreover, this value was higher than that obtained using DNA alignments. While DNA indel scores increased from A- to G-rich motifs, AA indel scores lack this pattern. It is important to note that C- and T-rich motifs exhibited a marked indel rate increase at the AA level, with median values of 0.45 and 0.25, respectively. G-rich motifs indel rates were different than T-rich motifs, but not than HS-PQSs (Pairwise Wilcoxon’s Rank Sum Test with P value adjustment for multiple comparisons, *P* = 5.5e^−12^ *and 0.22*). Thus, tandem repeats seem to promote higher rates of indel mutagenesis within coding regions, but the highest mutagenic effect corresponds to G-rich motifs potentially folded into very stable G4s.

**Fig. 6.**
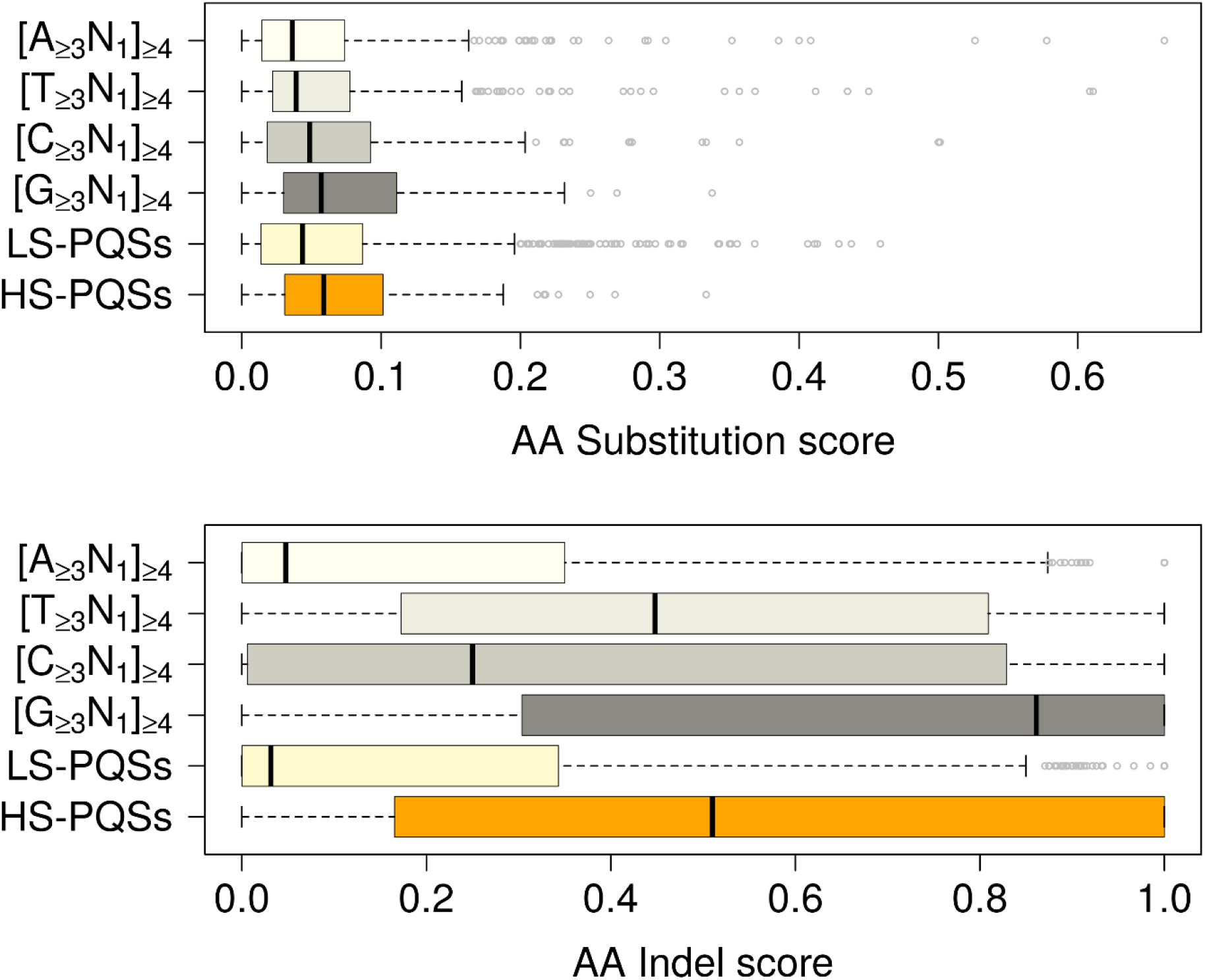
Amino acid mutational rates for PQSs and tandem repeat regions. **A** Substitution score comparison show that all analyzed motifs exhibited low lack of AA similarity across orthologous genes. **B** Indel rates comparison show that higher indel rates are associated to T-rich repeat motifs and structurally stable G4s.

A human-based GO-Biological processes analysis of genes containing HS-PQSs (n = 240) and LS-PQSs (n =1812) revealed significant overrepresentation of terms associated to developmental processes for both groups (Fig. 7). Thus, G4 occurrence in coding regions could be biased to genes involved in primate developmental regulation.

**Fig. 7.**
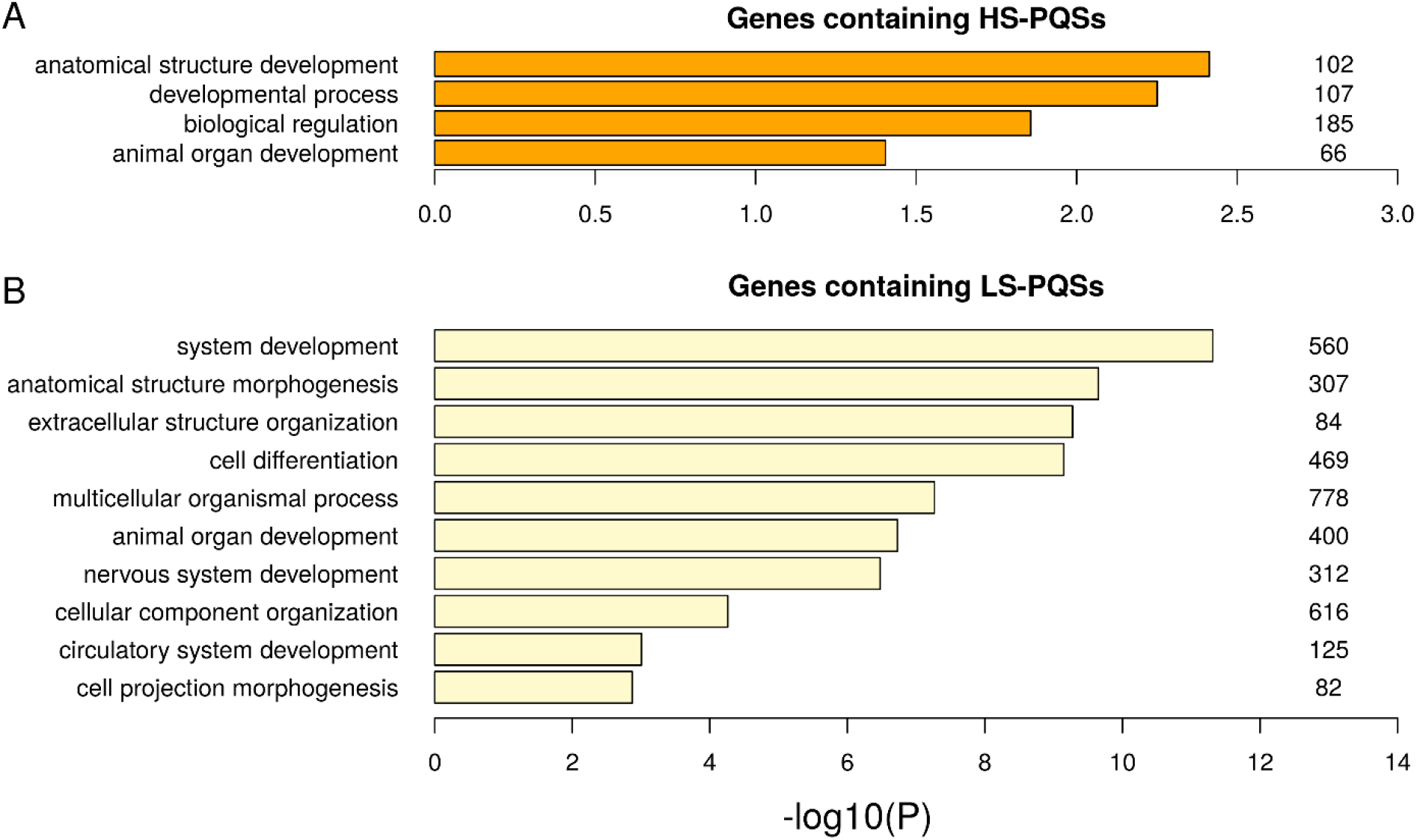
Gene overrepresentation analysis of genes containing HS-PQSs (**A**) and LS-PQSs (**B**) in terms of Biological processes. Gene count for each GO term is indicated to the right of each bar.

## DISCUSSION

Comparative genome analysis has demonstrated the high degree of protein coding regions similarity among primates. Here we present evidence that G-rich tandem repeats potentially folded into G4 structures depict the least conserved coding regions among primates. The G4-driven mutagenic effect during DNA replication and transcription have been described (Kruisserbrink et al., 2008; Yadav et al., 2014; Lemmens et al., 2015; Wang and Vasquez, 2017; Lopez et al., 2017), and this may explain their underrepresentation within coding regions. G4s occurrence have been extensively studied in different species (Marsico et al., 2019) and mostly using human-centric conservation analysis (Frees et al., 2014). We faced this issue by using parallel and non-biased multi-species approach during the identification and analysis of coding G4 conservation in primate ortholog genes. We found that stable G4s populate several developmental genes and mark low conserved and indel-rich coding regions. Moreover, this association was primate-specific or in close-related species, suggesting potential restricted implications without affecting the main protein function. Future work will be needed to determine if the G4s identified are functionally involved in the genome evolution of primate taxa.

In this study, we mapped and characterize G4 motifs within all available primate protein coding genomes. Our findings revealed that PQS occurrence varies depending on the species analyzed. The combination of computation tools with massive high-throughput sequencing methods has allowed researchers not only to identify G4 motifs across multiple organisms’ genomes, but also to assess their frequencies and conservation at important regulatory regions (Marsico et al., 2019; Verma et al., 2008; Yadav et al., 2008; Eddy & Maizels, 2008). However, human-biased approaches are commonly used when G4 occurrence and location are compared. In order to reduce this trend, we implemented a method that combined multiple species alignment of orthologous genes followed by PQSs search. For a proper G4 evolutionary analysis we focused on primate taxa. Interestingly, we were able to map efficiently both shared and species-specific PQSs. It is important to note that the genome assembly quality influenced the count of potential G4s finally identified, particularly mouse lemur and bushbaby. Consequently, PQSs were better characterized in hominid-related species. Thus, further efforts are necessary to clarify the coding G4 landscape in non-human primates.

Using a comparative primate genomics approach, here we show that very stable G4 motifs (G3N1)4 are underrepresented within coding regions and depict low conservation hotspots at the DNA and amino acid levels (Figs., 2 and 4). Very stable secondary structures, like G-quadruplexes, can interfere with translation efficiency in many RNAs (Bhattacharyya et al., 2014; Endoh et al., 2013a; Murat et al., 2014). Moreover, G4 are associated with high DNA mutational rates and genome instability (Lopes et al., 2011; Parekh et al., 2019; Wang et al., 2019). In accordance, recent studies have demonstrated that thermodynamically stable G4s are selected against in the coding region of mRNAs across species (Puig Lombardi et al., 2019; Mirihana Arachchilage et al..2019). Interestingly, the analysis of more than 500 species genomes revealed a strong accumulation of single-nucleotide-loop G4 motifs in primates, especially the thermodynamically least stable (G3A1)4 sequences. Therefore, the deleterious effects associated to the most stable G4s may explain their low frequency in coding regions, while the low conservation, but not complete deletion, associated to them suggest the existence of species-specific mutagenesis processes controlling the evolution of this motifs in primates.

RNA secondary structures can be positive and negatively selected depending on context (Katz and Burge, 2003; Shabalina et al., 2013; Gebert et al., 2019). Our results show that G4 stability influences their prevalence in coding regions. Highly conserved G4s with moderate/low stability may be preserved because of their potential roles on translation regulation of genes with similar expression patterns across primates (Endoh and Sugimoto, 2016;). While the deletion of more stable G4s may indicate an overall deleterious effect, stable G4s may be evolving to create more species-specific regulatory networks.

To gain more insights into the G4-associated mutagenesis pattern, we analyzed substitution and indel rates among potential non-B DNA repeats elements. Interestingly, G4 motifs seem to induce indel mutations, but not substitutions, at both, DNA and amino acid levels. A previous work reported that potential G4s are depleted of SNPs in humans (Nakken et al., 2009). More recently, Du et al showed that G4 structures are indeed associated with higher rates of mutagenic potential, although SNP or indel specific effect were not distinguished in this study (Du et al., 2014). One of the proposed justifications for the lack of concordance was the nature of used datasets. While Nakken et al. data had few indels (Sherry et al., 2001), that used for Du et al. (2014) included SNPs and indels (1000 genomes project, 2012). These results, although limited to human nucleotide variability, support our findings about the interspecies variability associated with G4s in coding regions.

DNA tandem repeats, including G4s, are mutagenic and expandable and have been implicated in nearly 30 human hereditary disorders, most of which primarily affect the nervous system through protein function alterations (Mirkin, 2007; Usdin, 2008; Paulson 2018). We argue that the low amino acid variability and high indel rates associated to G4s may be reflecting in-frame motif expansion events (López Castel et al., 2010; Renton et al., 2011; Iyer et al., 2015). Therefore, it is possible that stable G4s are likely arising by the expansion of G-rich tandem repeats in single or close-related primates. Another mechanism to explain coding G4 genome content is that ancestral G4 motifs are preserved only in specific species through functional selection. While G4-induced altered protein function could have deleterious effect for one species, as shown for humans (Fratta et al., 2012; Conlon et al., 2016), they could have a neutral/beneficial effect in other primates, depicting a mechanism of genetic diversification. The functional consequences of this potential G4-driven variability need further validation and could contribute to understand the effect of structure-prone sequences on the evolution of protein coding genome regions.

## MATERIAL AND METHODS

### Orthologous coding sequences (CDS) obtention and analysis

We downloaded CDS fasta sequences for all available primate species (n = 24) from Ensemble release 100 database (http://www.ensembl.org/info/data/ftp/index.html). The list with species scientific names, genomic data and PQS count is reported in table 1. For all species, original CDS fasta files were split and filtered to select the longest complete transcript (presence of a start codon ATG and a stop codon TAA/TGA/TAG) for each gene. Then, filtered human transcripts identifiers were used as input query for obtaining orthologs genes across primates using Biomart tool (https://m.ensembl.org/info/data/biomart/index.html). Ortholog genes lists were obtained separately for each specie and filtered to include nonhuman primate genes having a percentage of identity with those of humans equal or higher than 80%. Finally, for each gene having at least five orthologs across species, CDS sequences were joined into single fasta files (n = 17780).

### Multiple species alignment (MSA)

Ortholog gene CDSs were aligned using the msa R package (Bodenhofer et al., 2015); the alignment algorithm used was ClustalW with default parameters. In parallel, conservation scores and consensus sequences were calculated for each MSA using default parameters. Conservation scores were calculated using the *msaConservationScore()* method; in summary the method takes a MSA object computes the sum of pairwise scores for all positions of the alignment. Consensus sequences were obtained using the *msaConsensusSequence()* method, which takes a MSA object and computes a consensus sequence.

### Potential G-quadruplex forming sequences (PQS) identification

The quadruplex-forming G-rich sequence (QGRS) mapper algorithm was used to search for the sequence pattern 5′-G_x_L_≤7_G_x_L_≤7_G_x_L_≤7_G_x_-3′ where x ≥ 3 nucleotides and L represents the loops (Kikin et al., 2006). Each CDS within MSA was search for PQSs individually and based on PQS start and end positions we obtained unique motifs. This approach let us to identify all PQSs across species in an unbiased fashion, i.e. PQS can be present within either primate specie. Sequences located adjacent to the 5’ and 3’ PQS ends were defined as Upstream and Downstream PQS flanking regions. In principle, flanking regions should match PQS length, but if PQSs were located at the very start or end of CDS, flanking lengths could be shorter than PQS’s.

### Minimum free energy calculation

The minimum free energy (Mfe) was calculated for each PQS motif using the RNAfold tool from the ViennaRNA package (Lorenz et al., 2011). The option −g was used in order to incorporate the folding energy of G4 sequences. Then, PQSs were classified into two groups: i) High stability PQSs (HS-PQS) having Mfe value <= −30 kcal/mol and ii) Low stability PQSs (LS-PQS) having Mfe value >= −10 kcal/mol. The complete list, coordinates and Mfe of all mapped PQSs are reported in Supplementary Table S1. For each PQS, ten (10) random regions, matching PQS length and non-overlapping PQS positions in alignment, were obtained from consensus sequence and were split into high stability (HS-Rand) and low stability (LS-Rand) random groups using same criteria as for PQSs. These motifs were used as structural controls in our analysis.

### Tandem repeats search

Following the same approach for PQS identification, we used and in-house script to look for tandem repeat regions following the regular expression 5′-N_x_L_1_N_x_L_1_N_x_L_1_N_x_-3′, where N can be A,T,C or G, x ≥ 3 nucleotides and L represents single nucleotide loops. As for PQSs, only unique motifs were preserved.

### Motif conservation score calculation

Using the nucleotide conservation score vector obtained for each alignment, the conservation value of each PQSs, PQS flanking regions, HS-Rand, LS-Rand and tandem repeats were calculated. For that, conservation scores of each motif position within the alignment were obtained and the median of these values were calculated, depicting the motif conservation value. In addition, the average nucleotide conservation was calculated for all CDSs (CDS_ConScore_).

### Computation of Z-scores for all motifs

We calculated Z-score value for all motifs identified (PQS and non PQS) using the equation: (Motif_ConScore_ – mean(CDS_ConScore_))/sd(CDS_ConScore_).

### Nucleotide Substitution and Indel score calculation

We used the alignment consensus sequence, which can be a nucleotide or a gap, reference to calculate the substitution and indel scores per site. The substitution rate reflected the dissimilarity across species. We calculate indel, and not insertion or deletion, rates because when a *x* symbol (nt) in sequence *a* is aligned with a gap in sequence *b* (or vice versa), we have no way of knowing whether *x* was inserted in *a* or deleted from *b*. So, gaps were defined as indels.

If consensus was a nucleotide, we calculated: i) substitution score: all the nucleotides at the same alignment position was counted (nucleotide fraction) and the rate of dissimilarity with respect to the consensus was calculated by dividing the number of non-consensus nucleotides over nucleotide fraction, ii) indel score: all gaps (“-”) were counted and divide it by the aligned column size (count of aligned sequences).

If consensus was a gap, we estimated an overall indel score as 1 for that position.

Finally, all substitution and indel values of each motif position within the alignment were obtained and averaged, depicting the motif mutational rates.

### Amino acid Substitution and Indel score calculation

First, PQS and non-PQS motif coordinates were converted from DNA to AA. Then, motif-associated AA were aligned using an in-house script. We used a very similar approach to calculate amino acid substitution and indel scores. The unique difference was that we used as consensus sequence the most frequent symbol per site (AA or gap). The calculation was performed for each site and then averaged.

### Gene Ontology Analysis

Gene Ontology (GO) statistical overrepresentation analysis was performed with Panther classification system (Thomas et al., 2003). Functional categories potentially associated with genes containing PQSs were analyzed based on GOs biological processes and the significantly enriched categories were selected using an FDR threshold of 0.05.

### Phylogeny analysis

Mitochondrial RNA sequences of all placental mammals analyzed were obtained from Nucleotide database (https://www.ncbi.nlm.nih.gov/nuccore/) and aligned using the MAFFT alignment algorithm (Katoh & Standley, 2013). Finally, we obtained a phylo class object which was used for building phylogenetic tree using the *ape* (Paradis and Schliep, 2019) and *ggtree* (Yu, 2020) R packages.

### Statistical analysis

All statistical analysis was performed using the R software version 3.6.3 (R Core Team, 2020). The mutagenic rates variations were analyzed applying Pairwise Wilcoxon’s Rank Sum Test with P value adjustment.

## ACKNOWLEDGMENT

We acknowledge funding from the Brazilian National Council for Scientific and Technological Development (CNPq, grant 141004/2018-5 to MJE).

## AUTHOR CONTRIBUTIONS

MJE and SRP conceived and designed the work that led to the submission, acquired the data, interpreted the results and wrote the manuscript.

